# Study on the Difference of Prokaryotic Flora Structure on the Surface of Micro-nano Coating with Different Antifouling Property

**DOI:** 10.1101/435909

**Authors:** Qingze Gu, Aihan Zhang, Zhengyu Yan, Changlu Guo, Zhizhou Zhang

## Abstract

Low surface energy composite antifouling coatings prepared from carbon nanotubes (CNTs) and polydimethylsiloxane (PDMS) have good values for investigation of biofouling-related biological questions on marine biofilm. In order to deeply study the mechanism of antifouling on the surface of CNTs-PDMS coatings, it is necessary to investigate the structure of the microbial flora in the early biofilm on the coating surface. In the present study, the specific aim of this study was to investigate the structure differences of prokaryotic flora in biofilm samples at the early stage of biofouling through 16S rDNA based high-throughput DNA sequencing. By annotating high-throughput DNA sequencing results, this study identified dominant prokaryotic phyla and genera in biofilms of CNTs-PDMS coatings and identified significant differences in microbial composition and its dynamics among different coatings. Though the analysis of the Shannon index, Simpson index, Chao1 index and ACE index, coatings with better antifouling properties and antifouling properties have significant differences in community diversity and abundance, indicating different antifouling properties can affect the type and content of biofilm communities. According to the canonical correspondence analysis (CCA), time and temperature are more related to microbial community distribution, while the diameter and length of nanomaterials are less correlated. Through this study, the differences in microbial composition and content of prokaryotic communities, differences in diversity and abundance of sample communities, the differences between multiple samples and the correlation with important environmental factors were preliminarily analyzed, which laid a decent foundation for further research on the mechanism of anti-fouling on the surface of CNTs and PDMS coatings.

## INTRODUCTION

Marine biofouling refers to the general term for animals, plants and microorganisms attached to the surface of marine structures such as underwater artificial structures and ships, causing damage or adversely affecting human economic activities [1]. Antifouling coatings are widely used to remove the attachment or growth of fouling organisms. As an early antifouling coating, toxic compounds such as organotin, cuprous oxide and synthetic insecticides are commonly used as antifouling agents. However, antifouling coatings containing toxic compounds seriously damage the marine ecological balance and cause marine pollution problems, so the above antifouling agents have been banned or restricted [2,3]. Therefore, the development of non-toxic, long-lasting nanostructure-based coatings is one of the directions of future antifouling methods.

Low surface energy antifouling coatings with polydimethylsiloxane (PDMS) as the main raw material are the hotspots of marine antifouling coatings research, and the most promising antifouling coatings in the future marine antifouling field [4]. It utilizes the characteristics of low surface energy to reduce the adhesion efficiency of marine fouling organisms, making it difficult to adhere firmly to the surface of hull or marine building facilities, thereby inhibiting the growth of marine fouling organisms [5], though the already acquired antifouling capacity of PDMS is far away from satisfaction. PDMS coating is inefficient in both antifouling capacity and mechanical property. So its use in inhibiting marine biofouling is still highly limited. By introducing various biocides and nanofillers into the PDMS matrix, it is possible to improve its mechanical properties while keeping anti-fouling ability. Studies have shown that the incorporation of 0.1% (w/w) carbon nanotubes (CNTs) nanoparticles in the PDMS matrix can greatly enhance its ability to resist biofouling [6–12,19], but the authors did not find expected improvement in mechanical properties. Besides the endeavor direction of mechanical property augment, another direction is to modify surface biofilm dynamics and acquire better antifouling performance.

The formation of marine biofouling is an ecological succession process that includes several crucial stages, such as the adsorption of organic molecules, the accumulation of microorganisms such as bacteria and certain algae, and the attachment of larvae and algae spores of large fouling animals [13,14]. The aggregate formed by the adhesion of these microorganisms is called biofilm [15]. The biofilm formed on the surface of hull and marine facilities during early fouling has become a research hotspot of antifouling technology [16]. Because as the fouling flora formed in the early stage of marine fouling, biofilm’s composition of the prokaryotic and eukaryotic microorganisms directly relates to the early-stage attachment and later-stage growth of macrofoulers’ larvae [17]. Therefore, more and more scholars have begun to study the composition and structural characteristics of biofilms during the fouling process. Beigbeder et al [18] found that by adding multi-walled carbon nanotubes (MWNTS) to PDMS, the antifouling properties of PDMS coatings can be effectively enhanced. However, there have been lacking of studies on biofilm in the context of its compositional dynamics on the surfaces of nano-coatings prepared from carbon nanotubes (CNTs). Therefore, in order to deeply study the mechanism of antifouling on the surface of CNTs and PDMS coatings, it is necessary to investigate the structures of the microbial flora in the early biofilm.

In the present study, the antifouling properties of PDMS coatings with incorporated CNTs were examined. Biofilm samples were collected from these coating surfaces and then were subjected to microbial community structure analysis by next generation DNA sequencing. The specific aims of this study were: (1) to decipher the microbial compositions and observe their differential dynamic changes on the selected CNT-PDMS coating surfaces, and (2) to investigate the correlation relationship between the compositional structure of prokaryotic flora in the biofilm, marine environmental factor, and structural parameters of CNTs.

## MATERIALS AND METHODS

### Panel preparation

The steel test panels, measuring 100 mm × 100 mm × 3 mm, were thoroughly sanded with sandpapers in order to make the surfaces smooth. After polishing, carefully clean the surface of the steel panels with deionized water and 70 % (v/v) ethyl alcohol, and then rinse it repeatedly with deionized water for 3 times before drying it. Afterwards, these panels were coated with a layer of a non-toxic primer coat (i.e., the chlorinated rubber iron-red antirust paint) before experiments, so as to form a clear background for the outer layer, which is mainly made of a transparent silicone elastomer. In this study, the primer coat was purchased from HaoYing Company (Weihai, China) and it takes about 72 hours to cure at room temperature. It is worth mentioning that, according to previous research [19], the primer coating does not have the ability to resist biofouling.

### Preparation of PDMS composites

The silicone elastomer matrix used in this study was the Sylgard 184 kit (Dow Corning, USA), which was supplied as a two-part kit mainly consisting of a pre-polymer (base, part A) and a cross-linker (curing agent, part B). The six carbon nanotube materials used in the preparation of PDMS composites coatings were purchased from the Timesnano Company (Chengdu, China), including one single-walled carbon nanotubes (Short-SWNTs) and five MWNTs. The choice of carbon nanotube materials was related to the evaluation of the antifouling properties of CNTs-PDMS composite coatings. The specific evaluation criteria for the antifouling properties of different CNTs-PDMS composite coatings comes from the China national standard GB / T 5370 – 2007 — *Method for test of antifouling panels in shallow submergence*. According to the standard, the steel panels covered with antifouling paint were subjected to immersion test in shallow sea. After that, observe the variety of marine fouling organisms on the surface of the test panels, and evaluate and score the antifouling performance of the composite coating in accordance with the amount of bio-adhesion and the area covering the surface of the sample. Basing on the previous scoring results of anti-fouling properties of different CNTs-PDMS composite coatings in the laboratory, three CNTs-PDMS composite coatings with good marine antifouling properties and three CNTs-PDMS composites with poor marine antifouling properties were selected. Those PDMS composites were used as coating materials for the steel panels (the specific information of the CNTs fillers were shown in Table 1). When CNTs-PDMS coating is prepared, one side of each steel plate (already with primer coat as mentioned above) consumes 3 mL of A reagent, 0.3 mL of B reagent, and mg of nanomaterial. The above materials were mixed for 15-20 min before coating the test steel panels. After curing in an oven at 105°C for 2 hr and then 55°C overnight, the steel panels were ready for the subsequent field assays.

**Table 1.** The specific information of the CNTs fillers and PDMS composites

| Nanometer material | CNTs-fillers | OD (nm) | Length (μm) | Anti-fouling score<sup>[46-49]</sup> | Remarks |
| --- | --- | --- | --- | --- | --- |
| PDMS | -- | -- | -- | 61 | Control group |
| D12 | Carboxyl modified Short-SWNT2 | 1~2 | 1~3 | 91 | Three coatings with the highest antifouling and scoring |
| D26 | Hydroxyl modified MWNT2 | 8~15 | ~50 | 89 |  |
| D29 | Hydroxyl modified MWNT5 | 30~50 | ~20 | 89 |  |
| D32 | Carboxyl modified MWNT2 | 8~15 | ~50 | 62 | Three coatings with the lowest antifouling and scoring |
| D58 | Graphitized MWNT3 | 30~50 | ~30 | 63 |  |
| D59 | Graphitized MWNT4 | > 50 | 10~20 | 65 |  |

### Marine field assays

The test panels were immersed under the static conditions for 24 days at the XiaoShi Island harbor waters (N 37°31′51″; E 121°58′19″) locating on the northwest of Weihai, China. For each of the coatings, one panel was prepared in triplicate and immersed in 3 different locations in the same sea area. The steel panels were immersed in seawater at a depth of about 1.5 m, and taken out shortly for sampling at 3 d, 8 d, 15 d, and 24 d, respectively.

### Marine in situ experiment and sampling

The 24-day marine field experiment was conducted two times, from October 10th to November 13th and from November 11th to December 5th. The first batch of biofilm samples were collected on October 13, October 18, October 25, and November 3. The average seawater temperature during the sampling period was about 17.5°C (as shown in Table 1s). Before each sampling, the biofilm on the front and back surfaces of the steel plate was first photographed, and then the biofilm images of the coating surfaces were summarized and the differences were observed. The second batch of biofilm samples were collected on November 14, November 19, November 26, and December 5. And the average temperature of seawater during sampling was about 10.0 °C (as shown in Table 2s). Similarly, the biofilm on the surface of the steel sheet was photographed before each sampling. In the process of collecting biofilm samples in the early stage of fouling, disposable sterile gloves are required in order to avoid contamination of the sample by bacteria on the hands. First, the steel panels were taken out from the seawater, and the front and back sides of the steel panels were photographed. The steel panels were divided into 4 zones before sampling so that the effect of the previous biofilm sampling on the biofilm to be sampled can be reduced. The target area was first rinsed with sterile seawater, and a 1 mL pipette was used to add sterile sea water to the target area. Use a prepared small sterile brush to obtain the biofilm attached to the surface of the steel panels (larger fouling organism should be bypassed), and use a 200 μL pipette to aspirate the sterile seawater containing the biological sample and transfer it to the previously sterilized Eppendorf (EP) tube. After the sample was taken back to the laboratory, it was immediately centrifuged at 3000g for 5 min, and the obtained cell pellet was stored in a −80°C refrigerator for subsequent experiments. At the same time, according to the observed content of the biofilm sample, the amount of adhesion of the biofilm in each EP tube was counted, and a statistical table of the amount of adhesion was obtained. PDMS plus six CNTs-PDMS (total seven types of coatings) each had three steel panels with two side paints (so each coating had 6 sides). For sampling, a quarter of each side was sampled at each time point. So each coating was sampled 6 times at one time point. Two batches of field experiments collected 12 times of sampling for each coating.

**Table 2.** Biofilm samples and their groups

| Time | PDMS | PDMS +D12 | PDMS +D26 | PDMS +D29 | PDMS +D32 | PDMS +D58 | PDMS +D59 |
| --- | --- | --- | --- | --- | --- | --- | --- |
|  | RS1<br>(RS9) | RS2 | RS3 | RS4 | RS5 | RS6 | RS7 |
| DAY3 | RS1.1 | RS2.1 | RS3.1 | RS4.1 | RS5.1 | RS6.1 | RS7.1 |
|  |  | (RS10) |  |  | (RS11) |  |  |
| DAY8 | RS1.2 | RS2.2 | RS3.2 | RS4.2 | RS5.2 | RS6.2 | RS7.2 |
|  |  | (RS12) |  |  | (RS13) |  |  |
| DAY15 | RS1.3 | RS2.3 | RS3.3 | RS4.3 | RS5.3 | RS6.3 | RS7.3 |
|  |  | (RS14) |  |  | (RS15) |  |  |
| DAY24 | RS1.4 | RS2.4 | RS3.4 | RS4.4 | RS5.4 | RS6.4 | RS7.4 |
|  |  | (RS16) |  |  | (RS17) |  |  |

### High-throughput sequencing analysis

#### High-throughput sequencing

The metagenome DNA was extracted from a biofilm sample using Sangon Biotech (Shanghai, China) columnar soil genomic extraction kit (SK8263). The high-throughput sequencing of this 16S rDNA (V3+V4) region was based on the Ion S5 XL technology sequencing platform (Novogene Co. LTD, Beijing, China). The single-end sequencing (Single-End) method was used to sequence the 16S rDNA (V3+V4) region of 28 biofilm samples. The primer sequences used were 341F (5’-CCT AYG GGR BGC ASC AG-3’) and 806R (5’-GGA CTA CNN GGG TAT CTA AT-3’).

#### Species annotation and OTU analysis

Species annotation refers to the dominant species in terms of abundance at different classification levels (phylum, class, order, family, genus, specie) for each sequenced sample, and then using charting software to map the Relative abundance histogram, This makes it easy to observe the relatively abundant species and their total abundance at different levels of classification. To study the species composition of the early biofilm community, it is necessary to perform cluster analysis after processing the obtained sequence. The 97% similarity sequence is usually classified into the same class called Operational Taxonomic Units (OTUs), and subsequent compositional analysis is performed according to the representative sequence of the OTUs.

#### Sample complexity analysis

In order to study the diversity of sample microorganisms, one-sample diversity (Alpha diversity) analysis in community ecology can be used, and information such as community diversity and abundance can be obtained by this analysis. Two indices, the Chao1 index and the ACE index, are generally used to estimate the total number of species in the community. In order to characterize the abundance of the community in ecology, two indices, the Chao1 index and the ACE index, are generally used to estimate the total number of species in the community, which are represented by the number of OTUs in the community. The abundance is reflected by the size of the index. The larger the index, the higher the abundance of the representative community.

To characterize community diversity, two indices are introduced to predict community diversity and uniformity. Both the Shannon index and the Simpson index are used to predict community diversity. The former is more sensitive to rare OTUs, and the latter are more sensitive to dominant OTUs. They reflect the diversity of the community by the value. In general, the larger the Shannon index, the higher the diversity of the community. As for the Simpson index, it will have different values according to different algorithms. According to the algorithm of Qiime software used in this study, the larger the value, the higher the diversity of the community. In addition, in order to consider the credibility of the sequencing results, the sequencing depth index, Coverage, is needed to reflect the coverage of the sequencing. In general, the higher the Coverage value, the higher the probability that the sequence in the sample is detected, which means that the sequencing result can represent the actual situation of the sample, so that the subsequent analysis can be performed.

#### Analysis of species diversity curve

The dilution curve is a common curve that describes the sample diversity within a group. It can be used to determine whether the sequencing data is reasonable and to know the richness of the species in the sample. In the process of analysis, it is necessary to observe the trend of the dilution curve. When it tends to be flat, it indicates that the amount of sequencing data is reasonable, and more data will only produce a small number of new species.

The Rank-abundance curve is another way to analyze species diversity, which reflects the distribution of species (such as abundance and uniformity) from the OTU level. According to the relative abundance of OTUs sorted from large to small, the corresponding sort number is obtained. When drawing the Rank Abundance curve, the sort number is taken as the abscissa and the relative abundance is the ordinate. These points are connected in series by a curve, and the richness and uniformity information of the species in the sample can be obtained by observing the curve. The span of the curve in the horizontal direction represents the richness of the species, and the degree of tilt in the vertical direction represents the uniformity of the distribution. In general, the smaller the span of the curve on the horizontal axis, the smaller the species richness; the larger the span, the higher the species richness. And in the vertical direction, the smoother the curve, the more uniform the species distribution; the steeper the curve, indicating that the species distribution is more uneven.

#### Species accumulation boxplot analysis

The species accumulation boxplot is used to analyze the variation of species diversity with the increase of the number of samples. It is an effective tool for investigating the species composition of the sample and predicting the abundance of the species in the sample. In biodiversity and community surveys, the species accumulation boxplot is widely used to determine the adequacy of sample size and to estimate species richness. Through the species accumulation curve, it is not only possible to judge whether the sample size is sufficient, but also to predict the species richness under the premise of sufficient sample size. In order to examine the differences between groups, it can be analyzed by box plot, and the results visually reflect the median, dispersion, maximum, minimum, and outliers of species diversity within the group. At the same time, Wilcox rank sum test and Tukey test can be used to analyze whether the differences in species diversity between groups are significant.

#### Comparative analysis of multiple samples

Multi-sample comparative analysis is a comparative analysis of the microbial community composition of different samples. First, based on the species annotation results of all samples and the abundance information of OTUs, the OTUs information of the same classification is combined to obtain a species abundance information table. At the same time, the unisystem relationship between OTUs is utilized to further calculate the Unifrac distance. Unifrac distance is a way to calculate the distance between samples using evolutionary information between microbial sequences in each sample, and the distance matrix can be obtained if there are more than two samples. Then, using the abundance information of the OTUs, the Weighted Unifrac distance is further constructed based on the Unifrac distance. Finally, through the methods of Principal Component Analysis (PCA), Non-Metric Multi-Dimensional Scaling (NMDS), Unweighted Pair-group Method with Arithmetic Means (UPGMA) and Beta Diversity Index, the differences between samples are found.

#### Analysis of community diversity index between samples

In this work, the Beta diversity analysis is used to explore the heterogeneity of species composition between different habitat communities along the environmental gradient, and the similarities between different biofilm samples were obtained. To investigate the species with statistical differences between the groups, the LDA Effect Size (LEfSe) is performed in this study. LEfSe is an analysis tool for discovering and interpreting high-latitude data biomarkers. It can compare two or more groups to analyze the statistically different biomarkers between groups.

#### Principal component analysis

Principal Component Analysis (PCA) is a method of applying variance decomposition to reduce the dimensionality of multidimensional data to extract the most important elements and structures in the data. The PCA analysis can extract the two coordinate axes that reflect the difference between the samples to the greatest extent, so that the difference of the multi-dimensional data is displayed on the two-dimensional coordinate graph, thus revealing the simple law in the background of complex data. It is worth mentioning that the more similar the community composition of the samples, the closer they are in the PCA plot.

#### Non-metric multi-dimensional scaling analysis

Non-Metric Multi-Dimensional Scaling (NMDS) statistics is a sorting method suitable for ecological research. NMDS is a nonlinear model designed to overcome the shortcomings of linear models and better reflect the nonlinear structure of ecological data. The NMDS analysis reflects the species information contained in the sample in a multi-dimensional space in the form of points. And the degree of difference between different samples is reflected by the distance between points, which can reflect the differences between groups and within groups.

#### Unweighted pair-group method with arithmetic mean analysis

By constructing a cluster tree of samples, biofilm samples can be clustered to analyze similarities between different samples. Unweighted Pair-group Method with Arithmetic Mean (UPGMA) is a cluster analysis method often used in environmental biology research. The process of UPGMA analysis is to first gather the two samples with the smallest distance and form a new node whose branch point is located at 1/2 of the distance between the two samples. Then calculate the average distance between the new sample and other samples, and then find the smallest 2 samples for clustering. Repeat the above steps until all the samples are brought together to get a complete cluster tree. In the process of cluster analysis, UPGMA clustering analysis is performed with Weighted Unifrac distance matrix and Unweighted Unifrac distance matrix, and the clustering results are integrated with the species relative abundance of each sample at the door level.

#### Environmental Factor Association Analysis

In this experiment, the main environmental factors affecting the biome on the CNTs-PDMS coating include time (DAY), temperature (T), carbon nanotube material diameter (D), length (L), antifouling score (AF), and the degree of dispersion (DP) in PDMS. Canonical Correspondence Analysis (CCA) was used in this work to demonstrate the relationship between flora and environmental factors, which can detect the relationship between environmental factors, samples and flora and obtain important environmental drivers that affect sample distribution. When analyzing the CCA ranking graph, the environmental factors are represented by arrows in the graph, and the length represents the degree of correlation between the environmental factor and the community distribution and species distribution. So the longer the arrow length in the CCA sort map, the greater the correlation, and the shorter the arrow length, the less the correlation. The angle between the arrow line and the sorting axis represents the correlation between an environmental factor and the sorting axis. The smaller the angle, the higher the correlation; the lower the opposite. When the angle between the environmental factors is acute, there is a positive correlation between the two environmental factors, and a negative correlation when the angle is obtuse.

#### Next generation sequencing data availability

All original data in this article can be available upon request, and will be deposited in a public database before journal publication.

## RESULTS

### Marine field assays

In order to investigate the attachment of microorganisms, the amount of biofilm adhesion obtained per sample was counted in this study. According to the difference of biofilm adhesion in 1.5mL Eppendorf tube, a standard for semi-quantitatively judging the amount of biofilm adhesion is employed after sample centrifugation under 3000g for 3min, which is divided into 5 degrees of adhesion: (1) there is very little biofilm content in the tube; (2) the biofilm content in the tube just covers the bottom; (3) the biofilm content in the tube is about 2mm higher than the bottom; (4) the biofilm content in the tube is about 2~4mm higher than the bottom; (5) the biofilm content in the tube is about 4 mm higher than the bottom. The CNTs-PDMS coating surface adhesion data of the first marine field assays were processed to obtain an average value, which was used to draw a line chart, as shown in Figure 1s. It can be seen from that (1) the amount of adhesion of the biofilm increases with time; (2) the amount of biofilm adhesion on the CNTs-PDMS coating is generally higher than that of the PDMS coating; (3) among the composite materials, D26, D29, D58 and D59 have higher adhesion, and D12 and D32 have lower adhesion. The analysis of the adhesion of PDMS and CNTs-PDMS coatings showed that there was no obvious correspondence between the amount of biofilm adhesion and the antifouling properties.

### Sample grouping

Biofilm samples on PDMS-based coatings obtained from four sampling of marine field experiments were numbered accordingly as in Table 2. Taking the PDMS coating as an example, the biofilm samples on the 3rd, 8th, 15th, and 24th days were named RS1.1, RS1.2, RS1.3, and RS1.4. Similarly, for the CNTs-PDMS coating with No.12 carbon nanotube material (D12), the biofilm samples on the 3rd, 8th, 15th, and 24th days were named RS2.1, RS2.2, RS2.3, and RS2.4. Each sample was repeated for 12 times. So the total 336 biofilm samples were collected. Among them, 12 samples for each coating at the same sampling point were mixed into one sample, and there were 28 such mixed samples subjected to high-throughput DNA sequencing. In the case of high-throughput sequencing, in order to facilitate subsequent analysis of the data and to understand the differences between the biofilm sample sets, the sequenced biofilm samples need to be grouped. The first biofilm sample group was set according to the sampling time and the surface of the coating. The PDMS coatings, three CNTs-PDMS composite coatings (D12, D26, D29) with good antifouling performance and three CNTs-PDMS composite coatings (D32, D58, D59) with poor antifouling performance were set to 7 biofilm sample groups, respectively. The specific grouping settings are shown in Table 2. This grouping combines the sampling time and the anti-fouling performance, and set three kinds of carbon nanotube coatings (D12, D26, D29) with better antifouling performance and three kinds of carbon nanotube coatings (D32, D58, D59) with poor antifouling performance into different samples group. When performing high-throughput sequencing, the two grouping information is submitted to the sequencing company for analysis of the sequencing results. At the same time, bioinformatics methods were used to annotate and analyze the species composition and distribution of biomes in biofilm samples, and the differences between biofilm sample groups and environmental factors with large correlations with biome distribution.

### OTU analysis and species annotation

#### Statistics of species annotation results

As can be seen from Figure 2s, 28 biofilm mix samples were sequenced to obtain the corresponding number of Reads. Taking RS1.1 as an example, the abscissa in the figure represents the name of the biofilm sample, and the ordinate on the left represents the number of Reads. The red column height (Total Reads) represents the total number of Reads for the RS1.1 sample of 76,377 (i.e. effective data for subsequent analysis such as OTU clustering). The blue column height (Taxon Reads) represents the RS1.1 sample used to construct the OTUs and the number of Reads to obtain annotation information is 52,528. The tea green column height (Unclassified Reads) represents the number of Reads for which the RS1.1 sample does not receive annotation information is 4.The orange column height (Unique Reads) represents the number of reads in the RS1.1 sample that cannot be clustered into OTUs is 23,848 (i.e. sequences that cannot be clustered into OTUs will not be used for subsequent analysis). The ordinate on the right is the number of OTUs, so the purple column height (OTUs) represents the number of OTUs obtained by the RS1.1 sample is 1,024. Similarly, OTUs clustering and annotation statistics of other biofilm samples can be obtained.

#### Analysis of relative abundance of species

According to the results of the species annotation, the prokaryotic microorganisms of the biofilm samples have 39 phyla, 91 classes, 134 orders, 226 families, and 443 genera. The most dominant phyla are Proteobacteria, Cyanobacteria, Gracilibacteria and Bacteroidetes. The abundances of the top 10 phyla with the highest content in the biofilm samples were shown in Figure 1 below.

**Figure 1.**
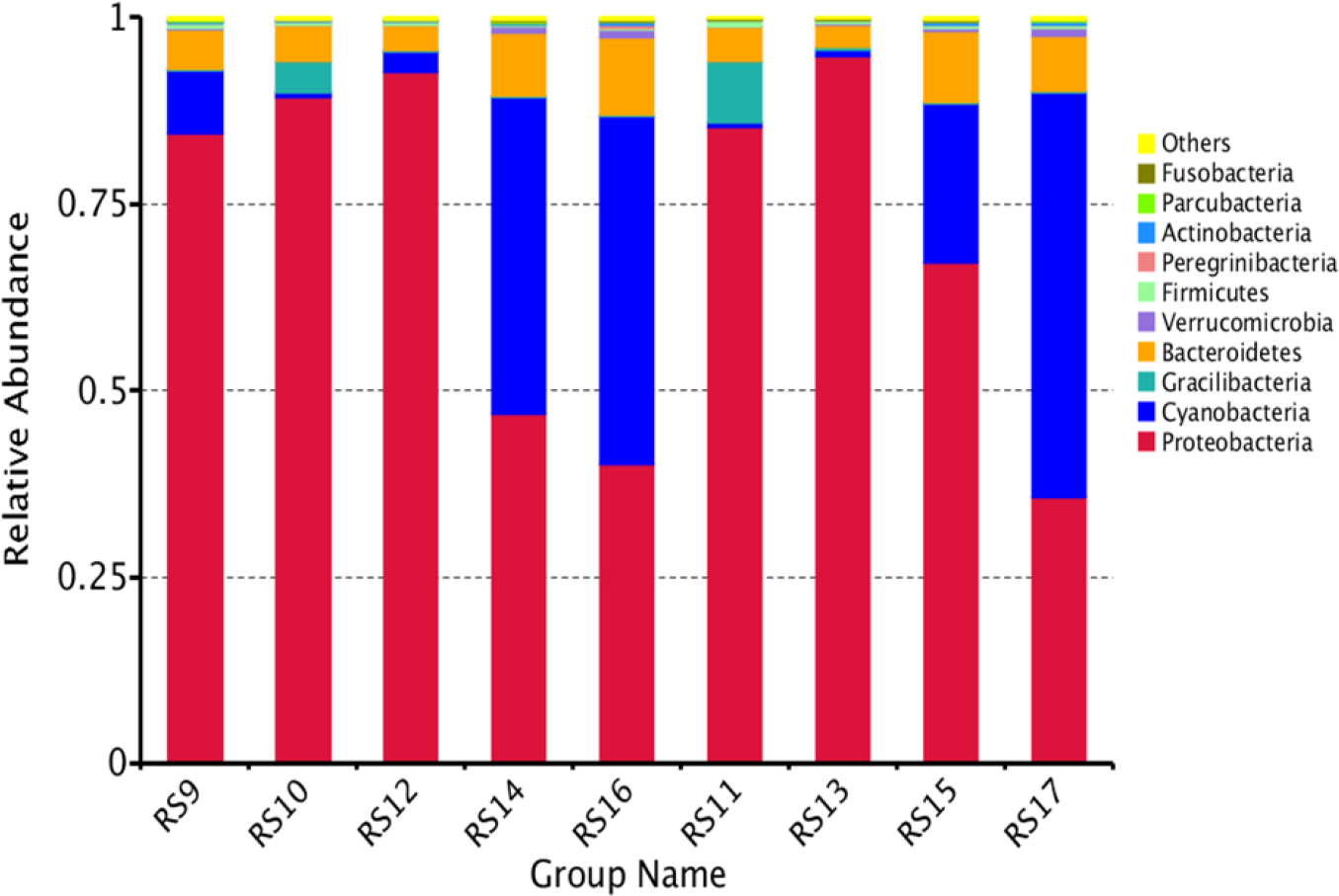
The relative abundance column of 10 dominant microbes at the phylum level

According to Figure 1, (1) the phylum with more content in the bacterial community of each group of biofilm samples is Proteobacteria. Among the four group samples RS10, RS12, RS14 and RS16 with better antifouling performance, Proteobacteria accounted for 89.40%, 92.80%, 46.80% and 40.21%, respectively. Among the four group samples RS11, RS13, RS15 and RS17 with poor antifouling performance, Proteobacteria accounted for 85.35%, 94.92%, 67.17% and 35.61%, respectively. It indicates that the Proteobacteria is the dominant phylum in the bacterial community regardless of the antifouling properties and time changes of the coating. (2) There were several other phyla with more content in the bacterial community of each group of biofilm samples, such as Cyanobacteria, Gracilibacteria and Bacteroidetes. (3) Regardless of the antifouling performance of the adhered coating, the content of cyanobacteria in the first two samples was less, accounting for 0.66%, 2.57%, 0.64%, and 0.78% in RS10, RS12, RS11, and RS13, respectively. However, the phylum cyanobacteria is more abundant in the last two sampling results, accounting for 42.61%, 46.67%, 21.40%, and 54.42% in RS14, RS16, RS15, and RS17, indicating that the content of cyanobacteria has gradually increased with time. (4) In each biofilm sample group, Gracilibacteria was more abundant in RS10 and RS11, which were 4.18% and 8.23%, respectively. It indicates that the content of Gracilibacteria decreased rapidly with time. (5) As the Figure 1 shows, it can be seen from the phylum level that no matter whether the antifouling performance of the coating is good or bad, the bacterial flora on the coating is similar in structure and the difference is not obvious at the same sampling time.

Similarly, the distribution of prokaryotic microorganisms at the genus level was obtained by annotation, and the top 10 genera were labeled. At the genus level, the dominant genera are *Cycloclasticus*, Unidentified Chloroplast, *Alteromonas*, *Oleiphilus*, *Oleibacter*, *Neptuniibacter*, *Thalassotalea*, *Colwellia*, *Oceaniserpentilla* and *Sulfitobacter* (Figure 2).

**Figure 2.**
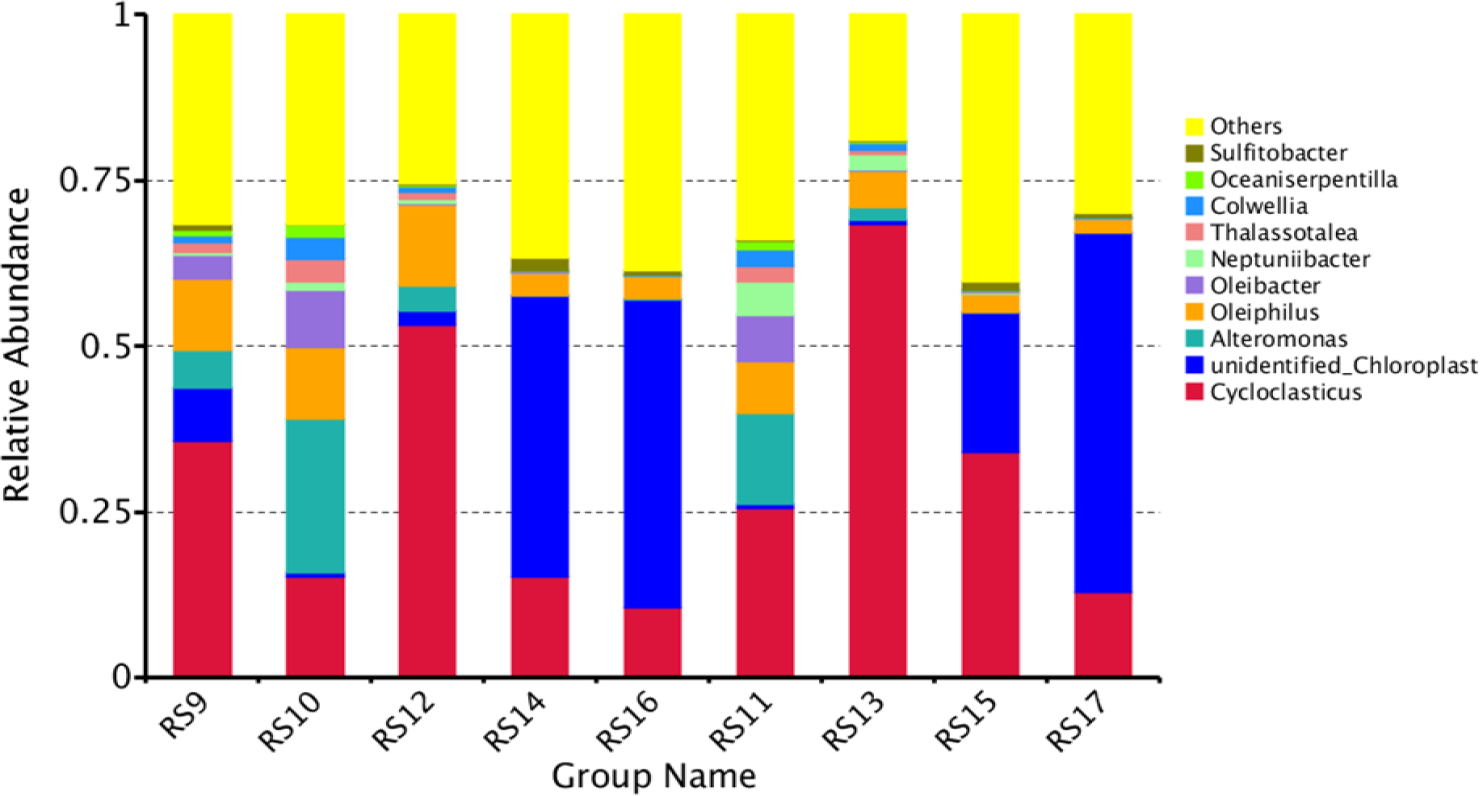
The spectral relative abundance column at the genus level

As can be seen from Figure 2, (1) in the bacterial community of each biofilm sample, *Cycloclasticus* accounted for 15.18%, 53.13%, 15.29% and 10.65% of the four biofilm samples RS10, RS12, RS14 and RS16 on the coatings with better antifouling performance, and accounted for 25.59%, 68.36%, 34.07%, and 12.99% of the four biofilm samples RS11, RS13, RS15, and RS17 on the coatings with poor antifouling performance. By comparison, it was found that *Cycloclasticus* contained more content on the coating with poor antifouling performance at the same sampling time point, and showed a trend of increasing first and then dropping down. (2) Among the biofilm sample bacterial communities in each group, Unidentified Chloroplast accounted for 0.58%, 2.253%, 42.33%, and 46.44% of the four biofilm samples RS10, RS12, RS14, and RS16 on the coatings with better antifouling performance, and accounted for 0.51%, 0.62%, 21.09%, and 54.15% of the four biofilm samples RS11, RS13, RS15, and RS17 on the coatings with poor antifouling performance. By comparison, it was found that Unidentified Chloroplast was gradually increasing with time. (3) The contents of *Alteromonas* in RS10, RS12, RS14 and RS16 were 23.25%, 3.84%, 0.04% and 0.06%, respectively, and the contents in RS11, RS13, RS15 and RS17 were 13.78%, 1.99%, 0.04% and 0.03%, respectively. It can be seen that *Alteromonas* exhibits a gradual decreasing trend on coatings with better and poor antifouling properties, and its content on the coating with better antifouling performance is higher. (4) The contents of *Oleiphilus* in RS10, RS12, RS14 and RS16 were 10.91%, 12.07%, 3.40% and 3.42%, respectively. It can be seen that *Oleiphilus* showed a tendency to increase first and then gradually decrease on the coatings with better antifouling performance. Also, it has a similar tendency to change in poor coatings. (5) In the bacterial community of each biofilm sample, *Cycloclasticus* appeared in large numbers. Studies have been conducted to show that *Cycloclasticus* may be destructive to antifouling coatings [20]. According to its function of degrading naphthalene and phenanthrene in seawater, this phenomenon may reflect the serious pollution of petroleum hydrocarbons near the sea area of XiaoShi Island harbor.

#### Alpha diversity index analysis

When studying microbial diversity in community ecology, the species abundance and diversity of microbial communities were understood through single-sample diversity analysis and statistical analysis. Table 3 below is a statistical table of the parameters of the microbial community alpha diversity of biofilm samples in the early stage of fouling. And the observed numbers of OTUs, sequencing coverage, and community diversity and abundance can be obtained from the data in the table.

**Table 3.**
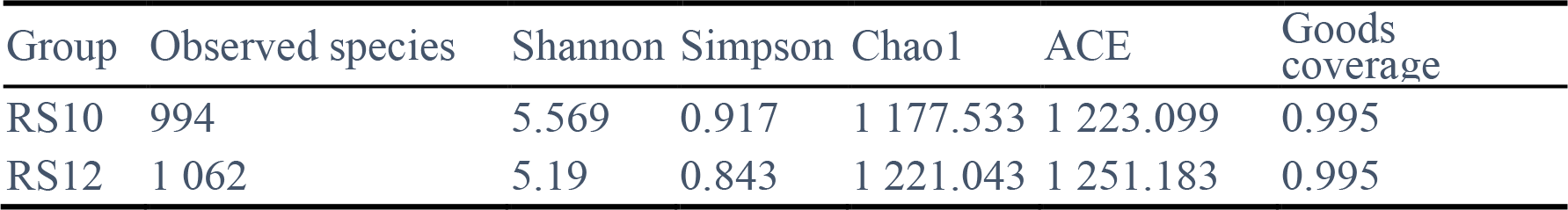
Statistics of microbial community diversity parameters of biofilm sample groups

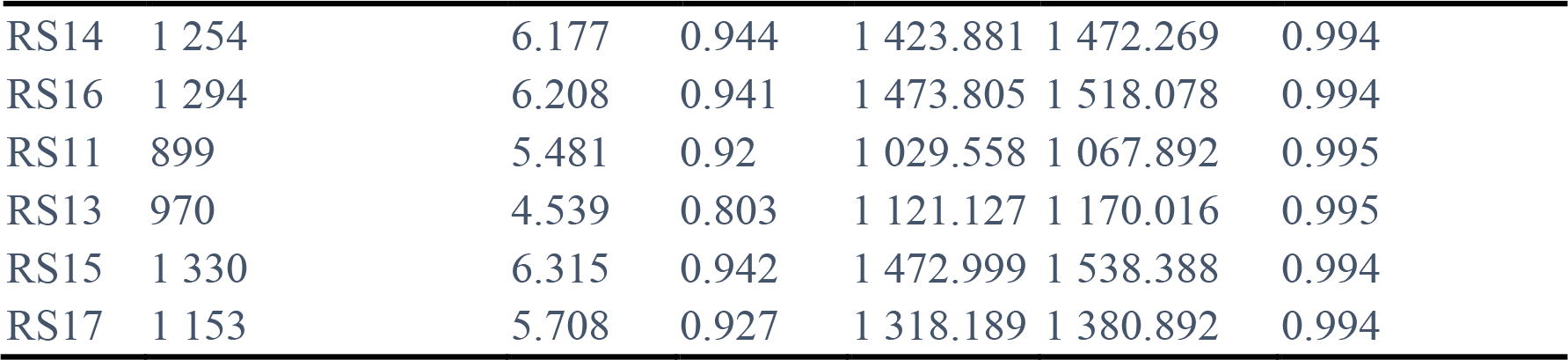

According to the data in Table 3, the following conclusions can be drawn: (1) The values of the Goods coverage of each biofilm sample group are between 0.994 and 0.995, indicating that the probability of the sequence being measured in each sample group is high and reflecting that the sequencing results can represent the true state of microorganisms in the biofilm sample group. (2) The comparison of the observed number of OTUs at the same sampling time point is: RS10>RS11, RS14>RS15, RS16>RS17, which suggests that the number of OTUs on the surface of the coating with better antifouling properties is generally greater than that of coatings with poor antifouling properties. It can be speculated that the diversity and abundance of the prokaryotic community of the coating with better antifouling performance is higher. (3) Comparing the number of observed OTUs at different sampling time points, it can be seen that with the increase of time, the biofilm sample groups tend to increase as a whole. And the longer the time is, the more diverse and abundant the biomes in the biofilm samples are. (4) By comparing the Shannon index and the Simpson index, it can be seen that the diversity of the prokaryotic community of the coating with better antifouling performance is basically higher. And the above conclusion is also confirmed by the comparison between the Chao1 index and the ACE index.

#### Species diversity curve

In order to further confirm the sequencing coverage of biofilm samples, it is possible to compare the richness of species in samples with different sequencing data by sparse graphs, and also to indicate whether the amount of sequencing data of the samples is reasonable. By analyzing the dilution curves of the 28 biofilm samples in Figure 3s, the biofilm samples and the curves of each group tend to be flat, indicating that the sequencing data is reasonable, and more data will only produce a small number of new species (OTUs). It can be clearly seen from Figure 3s that as the first and second sampled samples, the curves of the biofilm sample groups RS10, RS11, RS12 and RS13 tend to be flatter. For comparison, the curves of the sample groups RS14, RS15, RS16 and RS17 are relatively steep, indicating that the sequencing of the sample composition of the third and fourth sampling samples is greater than the previous two, and indirectly reflects the increase in the richness of species in the later samples. In order to understand the species abundance and species uniformity of the biofilm sample community, the Rank Abundance curve is needed to explain these two aspects of diversity.

By analyzing the Rank Abundance graphs of the 28 biofilm samples’ sequencing sequences in Figure 4s, it can be seen that the biofilm samples and the curves of each group tend to be gentle. In the horizontal direction, as the sampling time increases, the curve of the sample group RS10 to RS17 has a tendency to widen. The span of the curve on the horizontal axis becomes larger, indicating that the richness of the species in the later stage is higher. As the sampling time increases, the curve of the sample set RS10 to RS17 gradually becomes gentle from steepness, indicating that the uniformity of species in the sample gradually increases, and the distribution of species in the later period is more uniform.

#### Species accumulation boxplot analysis

Through the species accumulation boxplot, it is not only possible to judge whether the sample size is sufficient or not, but also to predict the species richness under the premise of sufficient sample size. By analyzing the species accumulation boxplot of the sequence of 28 biofilm samples in Figure 5s, as the sample size increases within a certain range, the position curve of the boxplot gradually becomes flat, indicating that even if the number of samples is increased, the species in the environment will not increase significantly. To conclude, the above analysis indicates that the biofilm sample size selected for this sequencing is sufficient and the sampling is sufficient for data analysis.

#### Difference analysis between groups of alpha diversity index

(1) Shannon index and Simpson index are used to understand the diversity and uniformity of the biofilm sample community. The diversity of microbial communities in biofilm samples in the early stages of fouling can be illustrated by the Shannon index and the Simpson index boxplots in Figure 3A.

**Figure 3.**
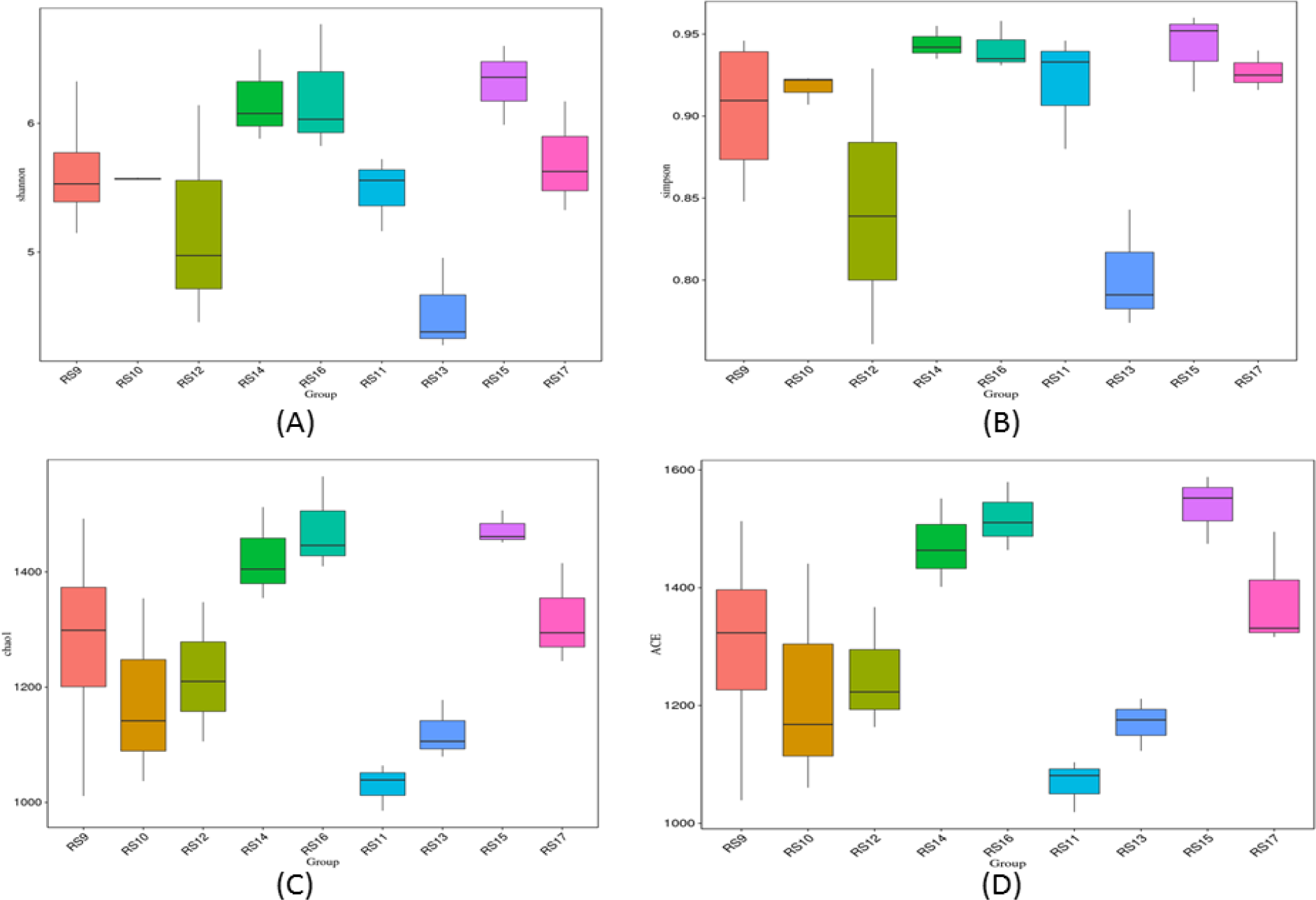
Plots of diversity indices for nine groups of samples. (A)The Shannon index; (B) The Simpson index**; (C)**The Chao1 index; (D)The ACE index.

There is a significant difference in the community diversity of the four different sampling times of the coating with higher anti-fouling score. Analysis of RS10, RS12, RS14, RS16 in the Shannon Wiener index and the Simpson index box plot shows that the community diversity obtained during the third and fourth sampling is higher, the diversity obtained during the first sampling was moderate, and the diversity obtained during the second sampling was low. Similarly, the community diversity of the coatings with poor antifouling scores at four different sampling times is significantly different, indicating that the prokaryotic microbial community diversity obtained during the second sampling is relatively low. Regardless of the Shannon index box plot or the Simpson box plot (Figure 3B), the analysis shows that the diversity of biofilm prokaryotic microbial communities on the coating with higher antifouling score has a trend of decreasing first and then increasing, while the change of the coating with poor anti-fouling score shows a trend of decreasing first, then increasing, and finally decreasing.

It can be seen from Figure 3A that the Shannon index of RS10 is not much different from that of RS11 at the same sampling time, but the Shannon index of RS12 is larger than RS13, RS14 is smaller than RS15, and RS16 is larger than RS17. According to Figure 3A, the Shannon index of RS10 at the same time point is smaller than RS11, but the Shannon index of RS12 is larger than RS13, RS14 is smaller than RS15, and RS16 is larger than RS17. It indicates that there is a difference in the prokaryotic community diversity of the biofilm samples on the coating with better and poor antifouling performance at the same sampling time. As can be seen from Figure 3A, there is a very significant difference between RS13 and RS14 (P=0.000 9<0.01), between RS13 and RS15 (P=0.000 4<0.01), between RS13 and RS16 (P=0.000 9<0.01), between RS13 and RS17 (P=0.016 6<0.05), indicating that there are differences in biofilm diversity at different sampling times, and some differences are significant.

(2) In order to investigate the abundance of the biofilm sample community (the number of OTUs in the community), the Chao1 index and the ACE index are required. By analyzing the Chao1 index and the ACE index boxplot of Figure 3C&3D, it can be seen that there is a significant difference in the abundance of the four different sampling times of the coating with higher antifouling score. According to the analysis of RS10, RS12, RS14 and RS16 by Chao1 index and ACE index boxplot, the microbial community abundance of biofilm samples gradually increased with the increase of sampling time. In addition, there is a significant difference in the diversity of four different sampling times for a coating with a poor antifouling score. Analysis of RS11, RS13, RS15, RS17 shows that with the increase of sampling time, the microbial community abundance of biofilm samples gradually increases and then decreases, but RS17 is still larger than RS11 and RS13, indicating that the microbial community abundance of biofilm samples is gradually increasing with time.

It can be seen from Figure 3C that the Chao1 index of RS10 is greater than RS11 at the same sampling time point, the Chao1 index of RS12 is larger than RS13, RS16 is larger than RS17, but RS14 is smaller than RS15. And It can be seen from Figure 3D that the ACE index of RS10 is greater than RS11 at the same time point, the ACE index of RS12 is greater than RS13, and RS16 is greater than RS17, but RS14 is smaller than RS15. It indicates that there is a difference in the prokaryotic community abundance of the biofilm samples on the coating with better and poor antifouling performance at the same sampling time.

It can be seen from Figure 3A that there are differences between the biofilm sample groups at different sampling time points, and some differences are significant. For example, there is a very significant difference between RS11 and RS14 (P=0.000 6<0.01), between RS11 and RS15 (P=0.000 1<0.01), between RS11 and RS16 (P=0.000 2<0.01), between RS11 and RS17 (P=0.011 6<0.05), indicating that there are differences in biofilm abundance at different sampling time points, and some differences are significant.

### Multi-sample comparison analysis

#### Analysis of differences between groups of beta diversity index

According to the boxplot, it is possible to analyze whether the differences in beta diversity between groups are significant. As can be seen from Figure 4, there is a difference in the beta diversity between the biofilm sample groups. And differences between biofilm samples exist regardless of the four different time points of the same coating or the same time point of the different coatings. However, there is no significant difference between the coating with better antifouling performance and the coating with poor antifouling performance at the same time point.

**Figure 4.**
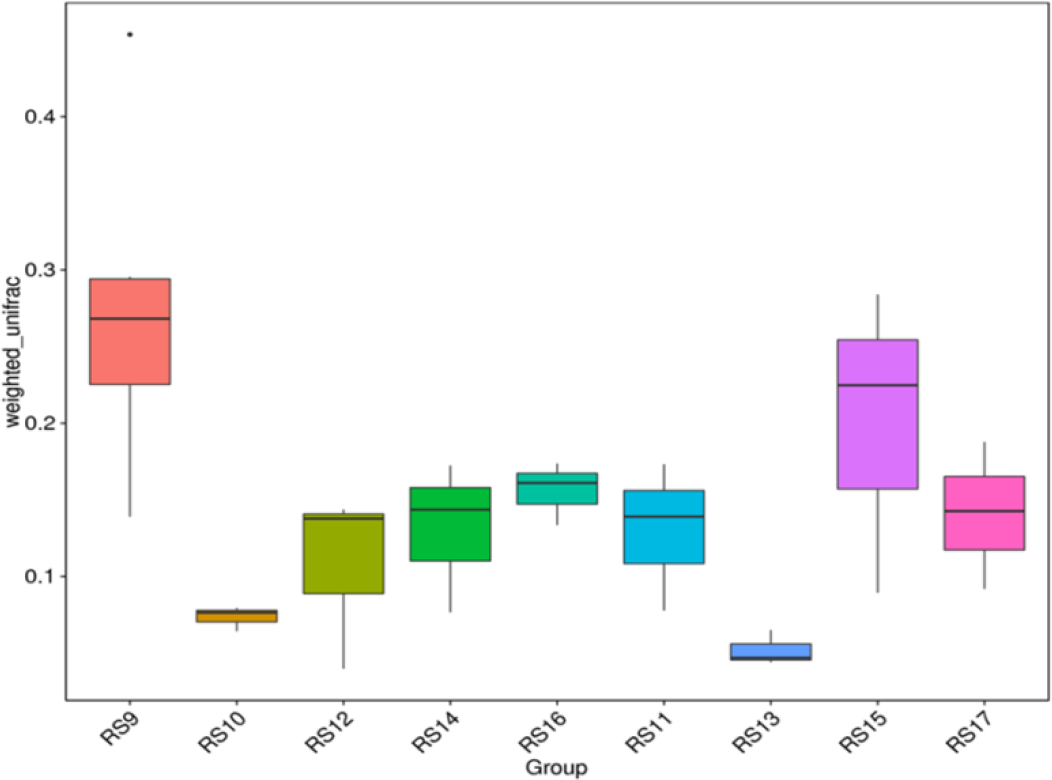
The beta diversity index boxplot for nine groups of samples

In order to analyze the statistically significant species between the biofilm sample groups, LEfSe was used to analyze the biofilm sample groups to obtain results at different taxonomic levels. Figure 5A shows the results of using LEfSe to analyze differences in the different taxonomic levels of biofilm samples from RS10, RS12, RS14, and RS16 in prokaryotic communities. The purpose of the analysis was to investigate species with significantly different antifouling properties for the three CNTs-PDMS coatings at four different sampling time points. The figure shows the significant difference species with the LDA score greater than the default value (the default value is set to 4.0), and the length of the histogram represents the LDA score, which is the degree of influence of significantly different species between different groups. Analysis of Figure 5A shows that RS10, RS12, RS14, and RS16 have distinct species at different taxonomic levels. Taking RS10 as an example, it is significantly different from the other three groups in phylum Gracilibacteria, class Gammaproteobacteria, order Alteromonadales, order Oceanospirilaceae, family Alteromonadaceae, family Oceanospirillaceae, family Colwelliaceae, genus Alteromonas, genus *Oleibacter*, genus *Thalassotalea*, and genus *Colwellia*. In addition, the analysis reveals that each of RS12, RS14, and RS16 was significantly different from the other groups. Similarly, analysis of Figure 5B shows that RS11, RS13, RS15, and RS17 have distinct species at different taxonomic levels. Taking RS17 as an example, it is significantly different from the other three groups in phylum Cyanobacteria, class Chloroplast, order Unidentified Chloroplast, family Unidentified Chloroplast, genus Unidentified Chloroplast, and species Virgulinella fragilis. And it can be seen that RS11, RS13, and RS15 each have a significantly different species relative to other groups.

**Figure 5.**
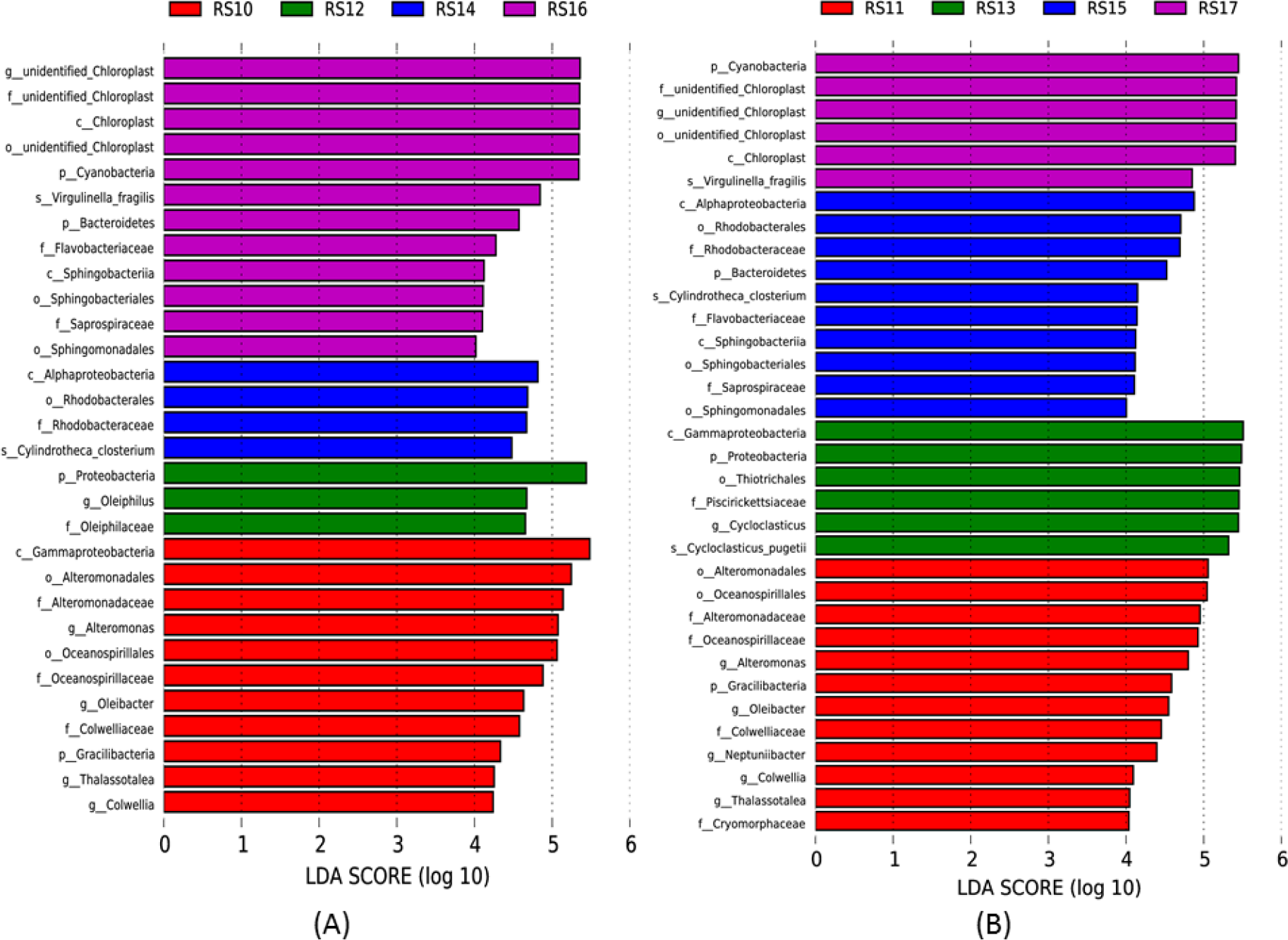
The LDA value distribution histogram of four biofilm samples on a coating with better antifouling performance (A) and four biofilm samples with poor antifouling performance (B).

#### Principal component analysis (PCA)

PCA is a method of applying variance decomposition to reduce the dimensionality of multidimensional data to extract the most important elements and structures in the data. If the community composition of the samples is more similar, the closer they are in the PCA plot.

From the information in Figure 6, we can see that: (1) At the same sampling time point, the components in the biofilm sample group are gathered together, and the points representing them are relatively concentrated, indicating that the difference between them is small. At different sampling time points, the components in the biofilm samples are far away, and their points are relatively dispersed, indicating that the difference between them is large and the time has a great influence on the biofilm samples. The above conclusions are consistent with the law of biofilm succession. (2) It can be seen from the figure that the points of RS10 and RS11 are more closely aggregated, indicating that the similarity between biofilm sample groups is very high in the early stage of fouling. However, the points of RS12 and RS13 are not very tightly aggregated. Similarly, the points of RS14 and RS15 and the points of RS17 of RS16 are not tightly aggregated, indicating that the similarity between biofilm sample groups gradually decreases with time and the difference between them begins to appear.

**Figure 6.**
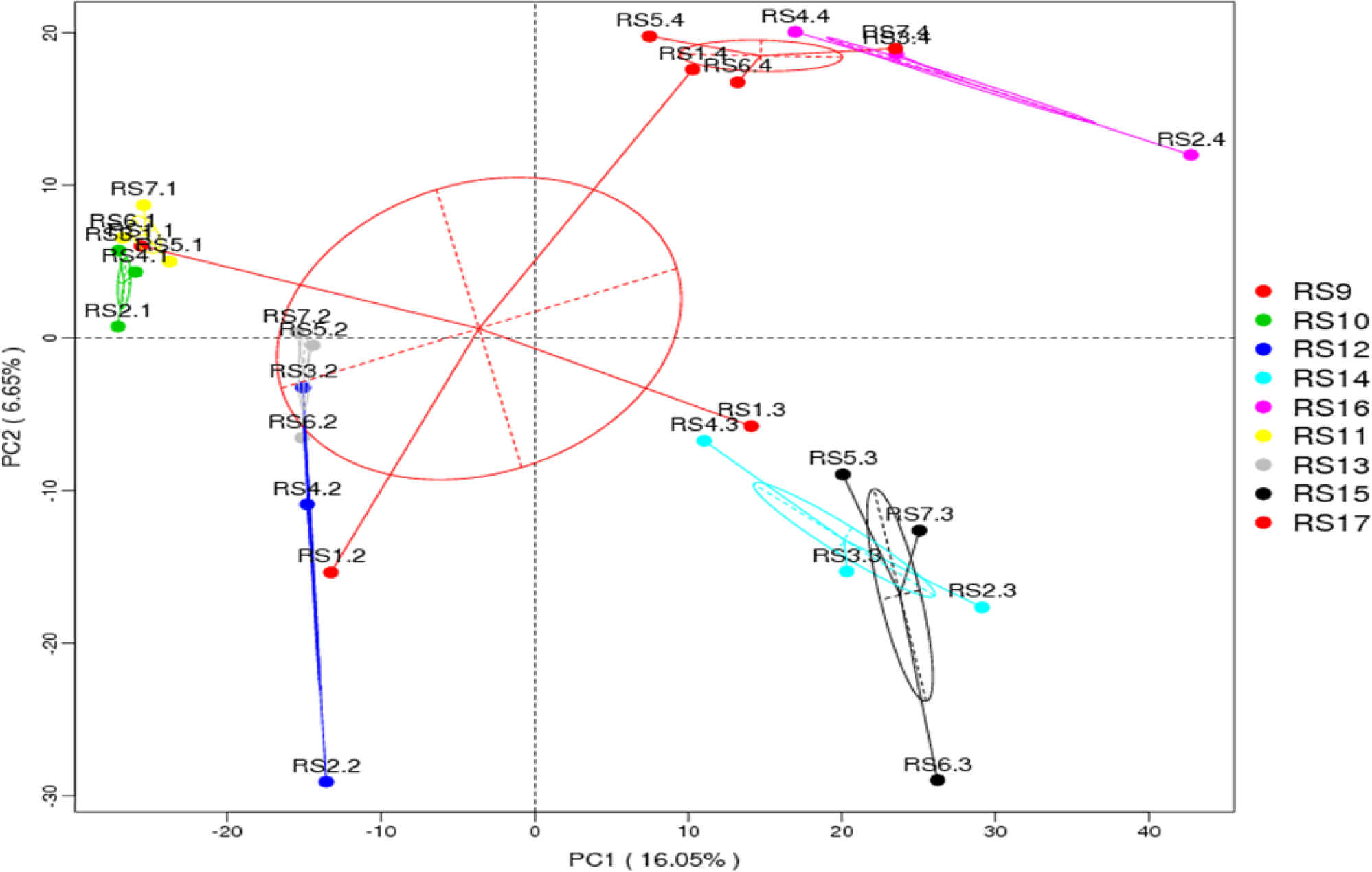
The principal component analysis chart of nine groups of high-throughput sequencing data

#### Non-metric multi-dimensional scaling (NMDS) analysis

NMDS is a sorting method for ecological research. Applying NMDS analysis can reflect differences between groups and within groups.

From the information in Figure 7, we can see that: (1) At the same sampling time point, the components in the biofilm sample group are clustered together, and the points representing them are relatively concentrated, indicating that the difference between them is small. At different sampling time points, the components in the biofilm samples are far apart, and their points are relatively dispersed, indicating that the large differences between them. (2) It can be seen from the figure that the points of RS10 and RS11 are more closely aggregated, indicating that the similarity between biofilm sample groups is very high in the early stage of fouling. However, the points of RS12 and RS13 are not very tightly aggregated, and the points of RS14 and RS15 and the points of RS17 of RS16 are also not tightly aggregated, indicating that the similarity between biofilm sample groups gradually decreases with time and the differences between them begin to appear. (3) By comparing with PCA, the degree of difference between each biofilm sample group can be known. And the value of Stress is 0.049 (< 0.2), indicating that NMDS can accurately reflect the difference between samples. (4) Compared with other biofilm sample groups, the points of RS14 are more concentrated, indicating that the similarity between them is extremely high, and the biofilm community had the smallest difference at that time. Similarly, RS15 had small differences at that time.

**Figure 7.**
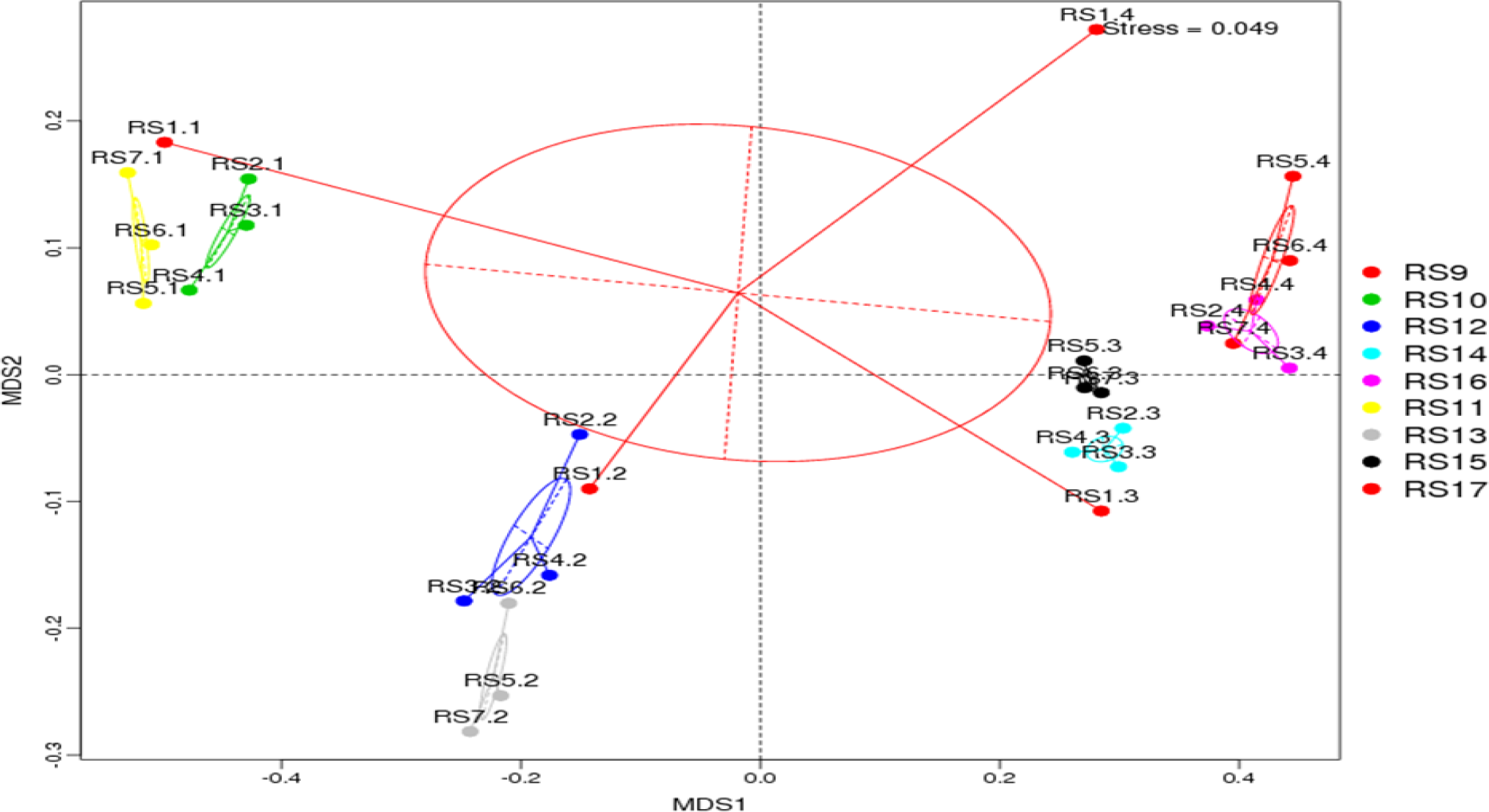
The NMDS chart of nine groups of high-throughput sequencing data

#### UPGMA clustering tree analysis

By clustering the samples and constructing a cluster tree, the similarity between different samples can be studied. According to the analysis in Figure 8, if the abundance of each species in the community is not considered to be different or the size of the difference, RS16 and RS17 have similar similarities and they are similar to RS14 and RS15. The same is true for RS12 and RS13, but the difference between RS11 and each group is larger and the similarity is lower. Through quantitative analysis of the UPGMA clustering tree, if the abundance of each species in the community is considered to be different or the size of the difference, the similarity between RS10 and RS11, between RS12 and RS13, and between RS16 and RS14 is higher. In addition, the similarity between RS15 and 14 is lower. It illustrates that the similarity between the coatings with better and poor antifouling performance is higher in the early stage of fouling, but the difference between them increases with time.

**Figure 8.**
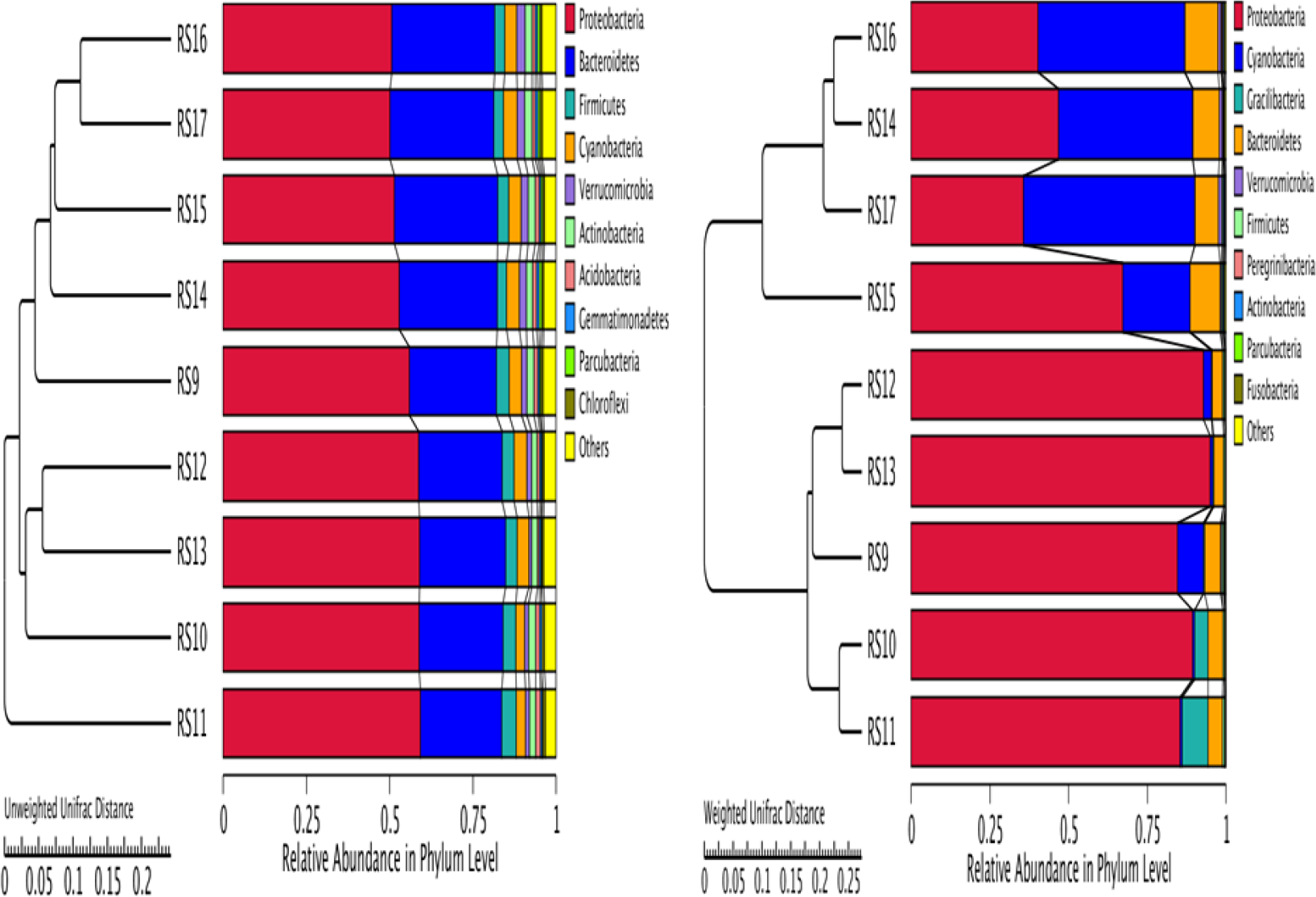
The unweighted (A) and weighted (B) UPGMA clustering tree for nine biofilm samples

#### Environmental factor association analysis

In order to study the effects of environmental factors on the composition and distribution of biofilm samples in the early stages of fouling, canonical correspondence analysis (CCA) is needed to analyze the relationship between biofilm flora and environmental factors, to identify the relationship between biofilm samples, biofilm communities and environmental factors, and to understand environmental factors that have crucial impacts.

As can be seen from Figure 9, (1) among the many environmental factors, the length of the arrow line of the sampling time DAY is the longest, indicating that the sampling time has the greatest correlation with the community distribution and species distribution. The length of the arrow lines such as AF and T is short, indicating that the correlation between antifouling performance, temperature and community distribution and species distribution is small. (2) According to the direction of the arrow of DAY, it indicates that the correlation between DAY and community distribution and species distribution in the third and fourth biofilm sample groups increases with the increase of sampling time. Similarly, the correlation between DAY and community distribution and species distribution in the first and second biofilm sample groups is smaller. (3) According to the direction of the arrow of DAY, it indicates that Unidentified Chloroplast and *Sulfitobacter* are positively correlated with the sampling time as the sampling time increases, and the correlation is great. Similarly, *Oleibacter*, *Alteromonas*, *Oceaniserpentilla*, etc. are negatively correlated with sampling time, indicating that Unidentified Chloroplast and *Sulfitobacter* are more likely to be produced in microbial communities in the later stages of sampling.

**Figure 9.**
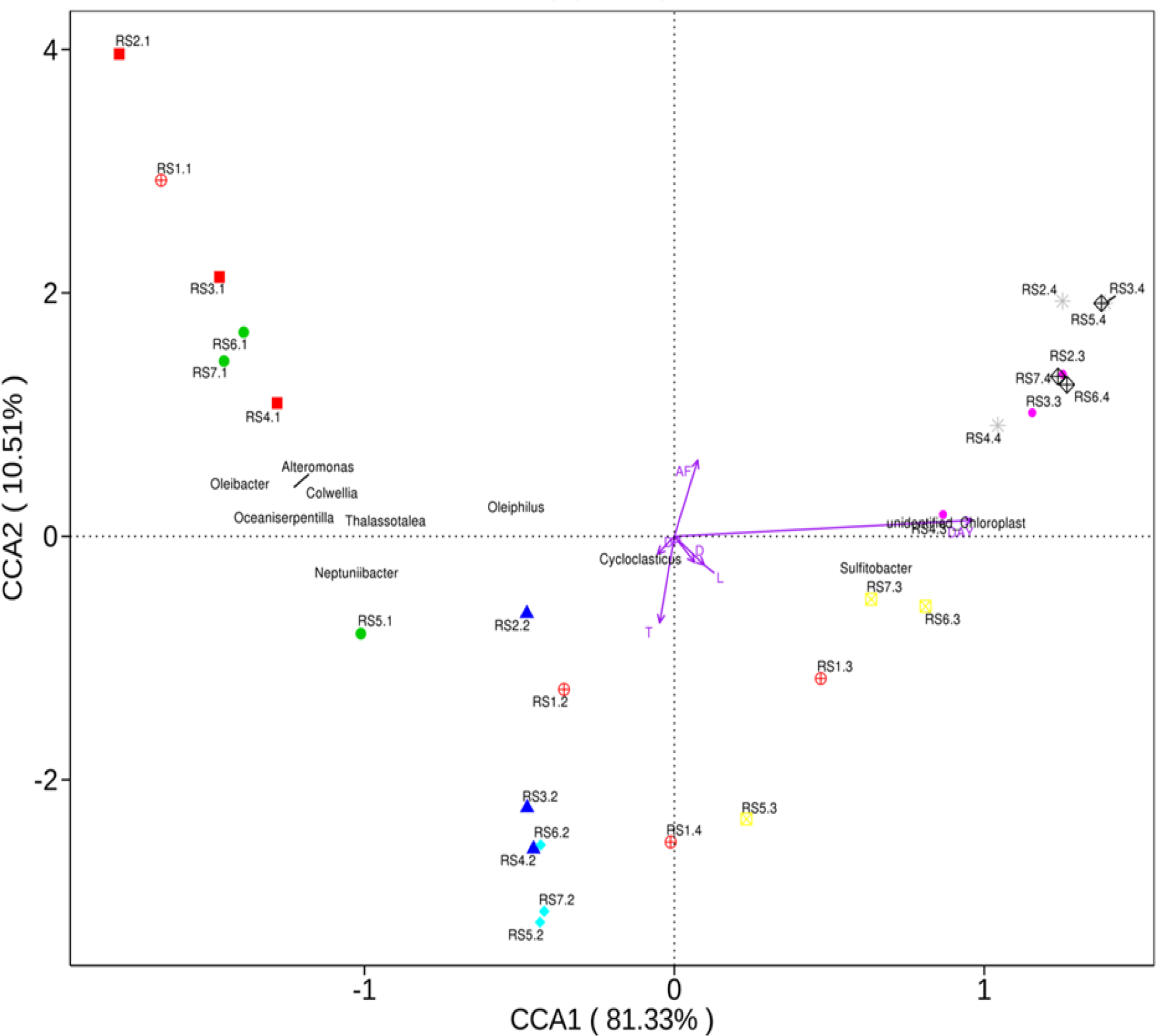
The sorting diagram of canonical correspondence analysis

## DISCUSSION

In the analysis of the species composition of the prokaryotic community, the dominant genus on the CNTs-PDMS composite coating was *Cycloclasticus*, *Alteromonas*, *Oleiphilus*, etc, which was different from the dominant bacteria on the surface of stainless steel, galvanized Q2358 and two marine paints [21–23]. The main dominant bacteria on the surface of reported stainless steel are *Pseudomonas* and *Pseudoalteromonas*. The dominant bacteria of galvanized Q2358 and titanium alloy are *Alteromonas* and *Pseudoalteromonas*. And the dominant bacteria on the surface of the two marine paints A and B are γ-Proteobacteria and α-Proteobacteria, respectively. It indicates that the microbial communities in the early stage of fouling on different materials have a large difference, or the surface properties of different materials have an effect on the composition of the bacterial community.

In this experiment, the prokaryotic microbial community has a large number of petroleum hydrocarbon-degrading bacteria such as *Cycloclasticus* and *Oleiphilus*, which are also present in two marine paints [23]. According to previous research, *Cycloclasticus* is not only a petroleum hydrocarbon degrading bacteria that can decompose naphthalene and phenanthrene, but also can cause damage to the coating. The large amount of petroleum hydrocarbon degrading bacteria also indicates that the seawater near the XiaoShi Island harbor in this marine experiment has been seriously polluted.

According to the analysis of community diversity, the coatings with better antifouling performance and the coatings with poor antifouling performance did not show significant differences. And the diversity and abundance of all prokaryotic communities increased, which showed the same trend as the diversity and abundance of prokaryotic communities on the coated surface of multi-walled carbon nanotubes (MWNTs) [23, 24]. It indicates that the surface of different CNTs-PDMS coatings does not cause a decrease in the diversity and abundance of the community or maintain a low level due to the poisoning effect on the antifoulant surface like the oxide of a heavy metal such as arsenic or copper [25, 26]. It also indicates that the antifouling mechanism of CNTs-PDMS coatings may be more complicated, perhaps relying on the interaction between community organisms, such as changes in content, or inhibition of metabolites, which requires more research to reveal.

For the analysis of environmental factors, the anti-fouling scores that were more concerned before the experiment had a small impact on the distribution of community and species, while time and temperature had a greater influence. According to the UPGMA clustering tree analysis, the coatings with better antifouling performance have greater similarity between RS14 and RS16 in the last two samplings. For coatings with poor antifouling properties, RS17 is similar to RS14 and RS16, while RS15 is less similar to them. It may be speculated that the biofilm on the coating with better antifouling performance may iterate faster and reach the “mature” state more quickly, while the biofilm renewal on the poor coating is iteratively slower. By changing the success rate of biomes, the dominant genus of biomes is more obvious, which affects the stability of the community, thus likely compromising the attachment efficiency of macrofoulers’ larvae. This speculation is similar to the results of Sun et al. [27], but more research is needed through subsequent experiments.

## CONCLUSION

The sample size of the biofilm selected by this sequencing is sufficient, the sampling is sufficient, the amount of sequencing data is reasonable, and the sequencing coverage is high. In addition, the sequencing results can represent the true state of the prokaryotic community in the biofilm sample groups and can be analyzed.

In the biofilm samples in the early stage of fouling, the dominant phyla are Proteobacteria, Cyanobacteria, Gracilibacteria and Bacteroidetes. Although the content of Cyanobacteria increased with time, the content of Gracilibacteria decreased rapidly with time, but the bacterial flora on the three coatings with better antifouling performance and three coatings with poor antifouling properties was similar in structure at the phylum level at the same sampling time, and the difference was not obvious.

Among the biofilm samples in the early stage of fouling, the dominant genus was *Cycloclasticus*, Unidentified Chloroplast, *Alteromonas*, *Oleiphilus*, *Oleibacter*, *Neptuniibacter*, *Thalassotalea*, *Colwellia*, *Oceaniserpentilla*, *Sulfitobacter*. With the increase of time, the content of Unidentified Chloroplast is gradually increasing, the content of *Alteromonas* is gradually decreasing, and the content of *Cycloclasticus* and *Oleiphilus* is increasing first and then decreasing. Compared with the level of the phylum, there is a significant difference in the genus level of the bacterial flora among sample groups.

According to the analysis of Shannon index and the Simpson index, regardless of whether the sampling time is the same, there is a difference between the coating with better antifouling performance and the coating with poor antifouling performance on the prokaryotic community diversity of the biofilm samples, and some differences are significant. For coatings with higher antifouling scores, the variation trend of the biofilm prokaryotic microbial community diversity is first reduced and then increased, while the coating change with poor antifouling score showed a trend of decreasing first, then increasing, and finally decreasing. It indicates that coatings with different antifouling properties have a certain influence on the diversity of prokaryotic communities.

According to the analysis of Chao1 index and ACE index, there is a difference between the coating with better antifouling performance and the coating with poor antifouling performance on the prokaryotic community abundance of the biofilm samples, regardless of the sampling time. And there are some significant differences. For biofilm samples with different sampling times, the abundance of microbial communities on coatings with higher antifouling scores and coatings with poor antifouling scores gradually increased, but the microbial community abundance on the coating with poor antifouling score first increased and then decreased.

It can be seen from PCA and NMDS results that the difference in composition in the biofilm sample group is small at the same sampling time point. At different sampling time points, the composition of the biofilm sample group is very different, and the time has a great influence on the biofilm sample, which is consistent with known facts in biofilm succession. As time increases, the similarity between the biofilm sample groups gradually decreases, and the difference begins to appear. In addition, the similarity of RS14 and RS15 in the biofilm sample group was extremely high, and the difference in biofilm community was the smallest.

Through the qualitative analysis of the UPGMA clustering tree, it can be seen that the similarity between the coating with good antifouling performance and the poor coating at the beginning of the fouling is high. But as time goes on, the difference between them gradually increases.

According to the CCA analysis, among the many environmental factors, the sampling time has the greatest correlation with the community distribution and species distribution, and the antifouling performance and temperature are ranked behind. Unidentified Chloroplast and *Sulfitobacter* are positively correlated with sampling time and are more likely to be produced in microbial communities in the later stages of sampling, while *Oleibacter*, *Alteromonas*, *Oceaniserpentilla*, etc. are negatively correlated with sampling time and are more likely to be produced in early sampling stages. It indicates that in environmental factors, time and temperature are highly correlated with community distribution and species distribution, while the diameter and length of nanomaterials are less correlated with community distribution and species distribution.

Through this study, the differences in species composition and content of prokaryotic communities, differences in diversity and abundance of sample communities, the differences between multiple samples and the analysis of important environmental factors were preliminarily analyzed, which laid a foundation for further research on the mechanism of marine anti-fouling on the surface of CNTs-filled PDMS coatings.

## ACKNOWLEDGEMENTS

This study was supported by the following funds: NSFC (No.31071170); GujingTribute fund (2016-1); GREDBIO (201401); the Key research and development plan of Shandong Province (2016GSF115022); and the Natural Science Foundation of Shandong Province (ZR2018MC002).

## Supporting documents

**Table 1s.** The first marine field experiment and sampling time

| Hanging panels and sampling | Hanging panels | 1<sup>st</sup> sampling | 2<sup>nd</sup> sampling | 3<sup>rd</sup> sampling | 4<sup>th</sup> sampling |
| --- | --- | --- | --- | --- | --- |
| Date | 2017.10.10 | 2017.10.13 | 2017.10.18 | 2017.10.25 | 2017.11.03 |
| Hanging time/day | 0 | 3 | 8 | 15 | 24 |
| Water temperature /°C | 20 | 17 | 18 | 20 | 15 |

**Table 2s.** The second marine field experiment and sampling time

| Hanging panels and sampling | Hanging panels | 1st sampling | 2nd sampling | 3rd sampling | 4th sampling |
| --- | --- | --- | --- | --- | --- |
| Date | 2017.11.11 | 2017.11.14 | 2017.11.19 | 2017.11.26 | 2017.12.05 |
| Hanging time/day | 0 | 3 | 8 | 15 | 24 |
| Water temperature /°C | 14 | 12 | 11 | 9 | 8 |

**Figure 1s.**
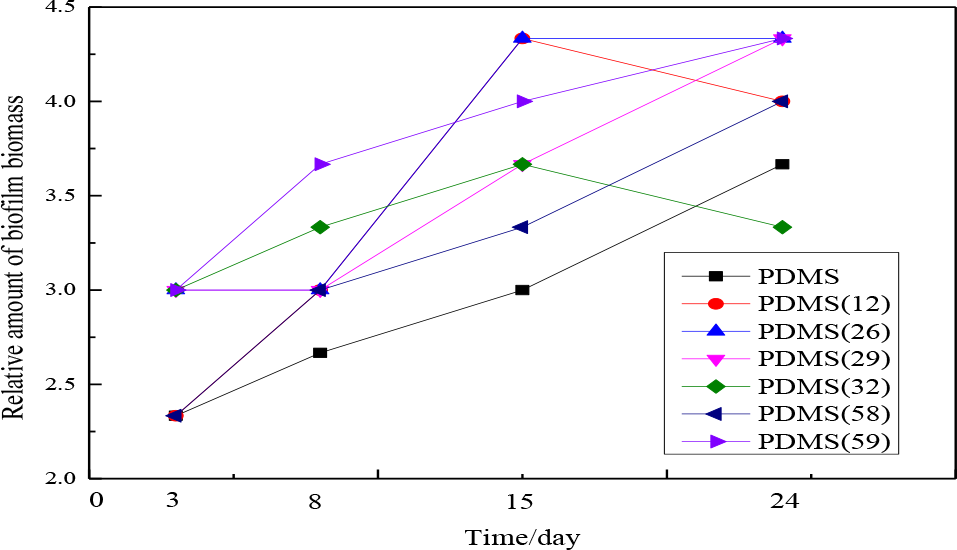
The relative biomass amount of biofilm gradually increased with the hanging time. Each point represented 3 replicates’ average value.

**Figure 2s.**
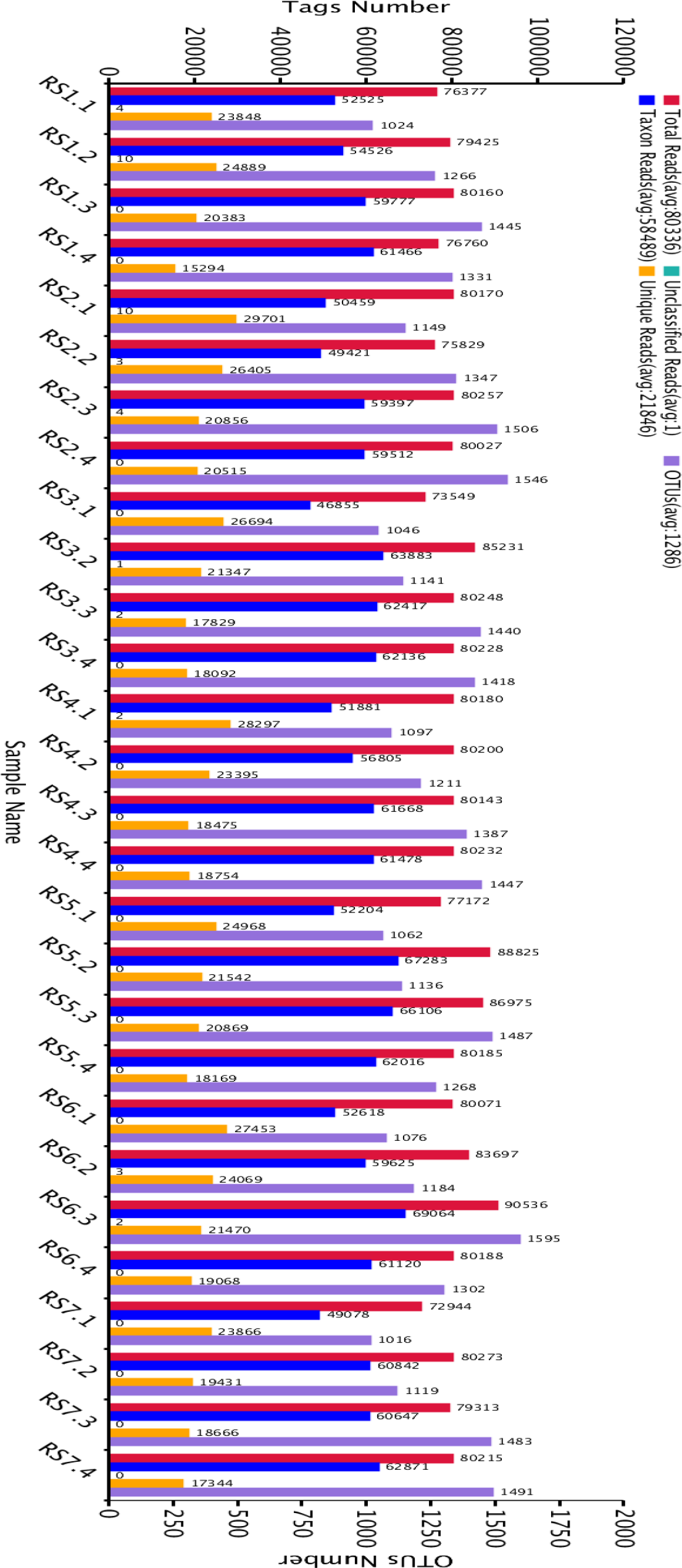
The OTUs clustering and annotation statistics for each biofilm sample

**Figure 3s.**
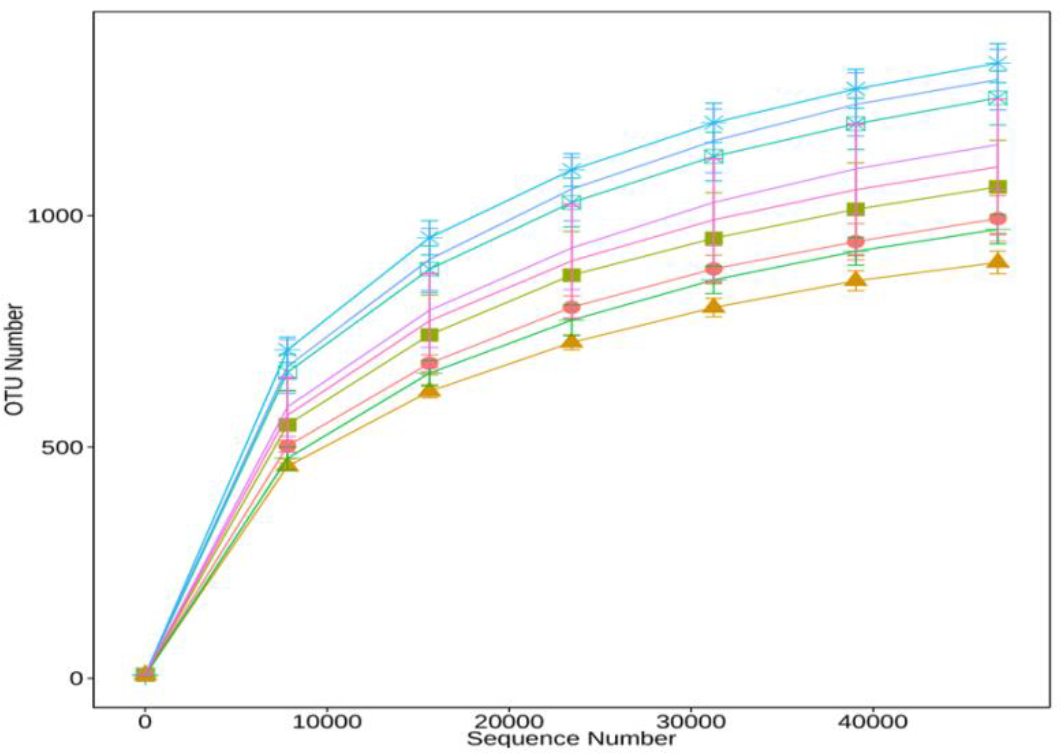
The biofilm sample group dilution curve

**Figure 4s.**
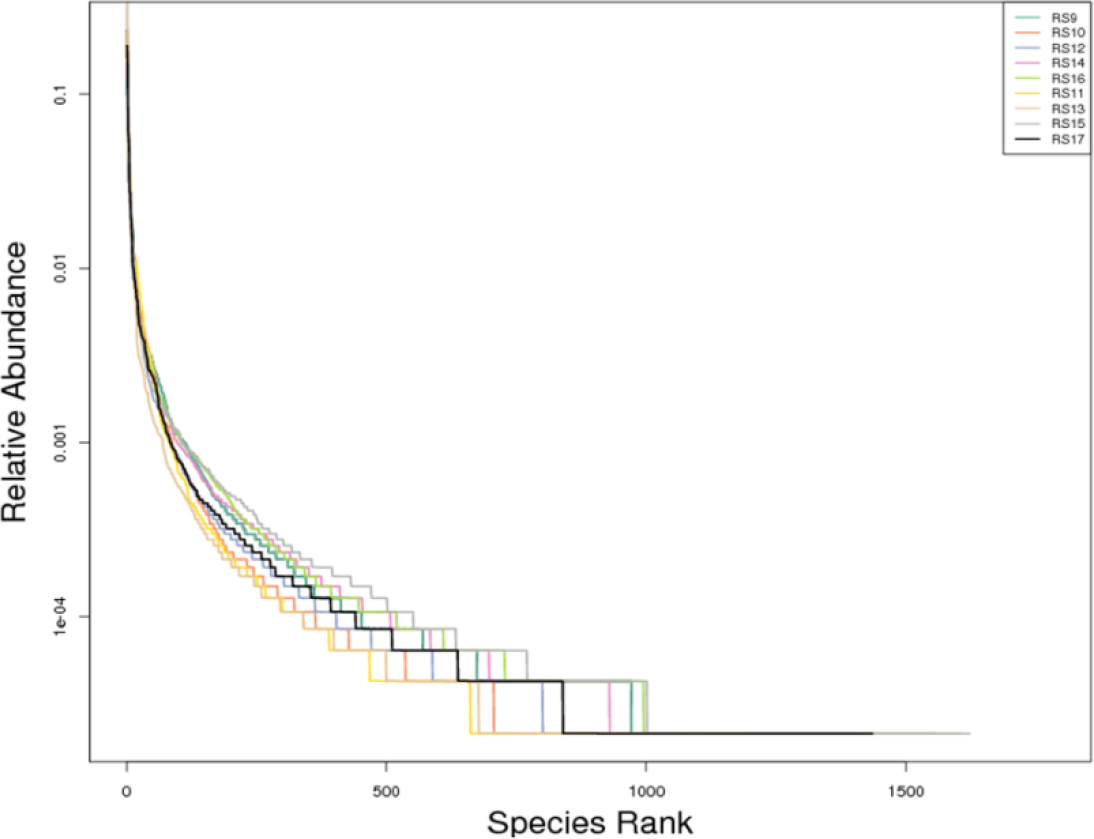
The rank Abundance chart of biofilm sample group

**Figure 5s.**
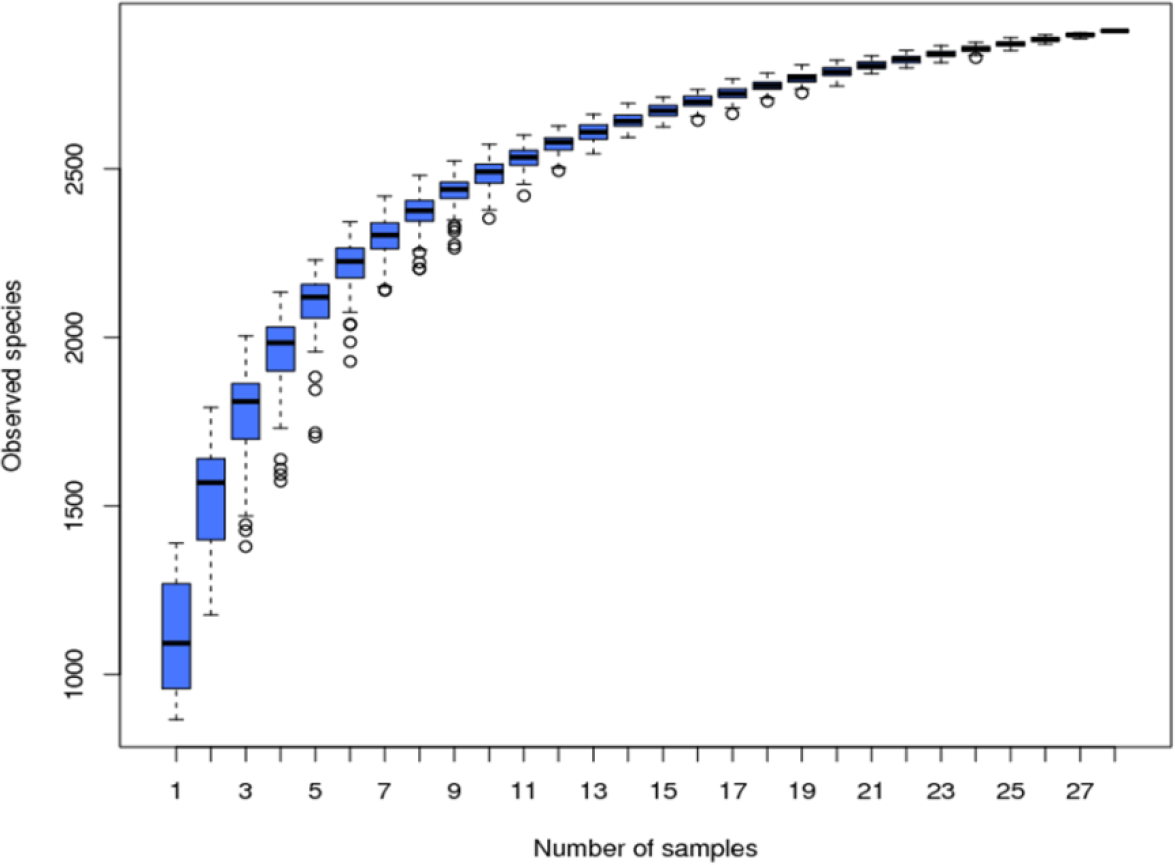
The species accumulation boxplot of 28 biofilm samples

